# Assessing the potential impact of Medicaid work requirements on African-Americans via a welfare reform analysis: a systematic review

**DOI:** 10.1101/549493

**Authors:** Garrett Hall, Sahai Burrowes

## Abstract

The Centers for Medicare and Medicaid Services are currently approving Medicaid demonstration projects involving work and community engagement requirements. The fear among public health professionals and policymakers is that these work requirements will disproportionately limit access to health insurance for African-Americans. To investigate these new work requirements for Medicaid, we conducted a systematic review of the Temporary Assistance for Needy Families program, the last major welfare program to have such work requirements. We used the ProQuest database and selected scholarly journal studies in English and published between January 1991-August 2018. Our selection strategy yielded 14 eligible studies, eight of which focused on caseload movements and six focused on sanctions. We found that African-Americans entered TANF at a higher rate than Whites, remained on TANF longer, and were subject to sanctions more frequently and stringently than Whites were. These results suggest that African-Americans may disproportionately experience reduced access to care through sanctioning such as lockout periods. Recommended policy changes include prohibiting work activities as a condition of Medicaid coverage, strengthening Medicaid in other ways, combating discriminatory hiring practices, and easing burdensome reporting requirements.

## Introduction

This paper investigates the potential impact of Medicaid work requirements on access to health insurance for African-Americans. We do this by conducting a systematic review of the literature on the impact of such work requirements on past government welfare programs.

Medicaid is a public health insurance program that covers 66 million low-income adults, children, pregnant people, disabled individuals, and the elderly [1]. Title XIX of the Social Security Act of 1965 created Medicaid along with Medicare under Title XVIII, which provides health insurance to the elderly population [2]. Medicaid provides health insurance coverage to low-income Americans based on applicants’ federal poverty level (FPL) and their eligibility within the categorical restrictions stated above. In 2017, for reference, the FPL for a family of four was an income of $24,600 [3]. Families earning this income are said to be at 100% of the FPL. The Affordable Care Act (ACA) enabled states to expand their Medicaid coverage to all adults at or below 138% of the FPL with no categorical restrictions [4]. Prior to the ACA, for example, categorical restrictions meant that low-income women must be pregnant or a parent to qualify for insurance through Medicaid; whereas, after the ACA and Medicaid expansion, any woman who meets the FPL requirement is able to enroll in the program [5]. Thirty-seven states and the District of Columbia chose to expand the program [6]. In a 2016 brief, the Urban Institute estimated that if the remaining 19 states decided to expand Medicaid, up to five million people would gain Medicaid coverage including 1.2 million African-Americans [4]. With states deciding to approve the program’s expansion and throw off the limiting categorical restrictions in the name of improving access, Medicaid began to live up to the ideals of its founding.

Medicaid has developed into a crucial part of the American safety net. It is the largest provider of public health insurance [7], insuring one-in-five Americans [8]. It is also the third most expensive federal domestic program and the federal government’s Federal Medical Assistance Percentage (FMAP) matching funds comprises the biggest federal contribution to state budgets [7,9]. Beyond its fiscal importance, Medicaid provides crucial health services to the marginalized. Slightly more than 40% of all Medicaid enrollees are children, while the elderly and people with disabilities account for approximately one in four enrollees [8]. A researcher of the social and political impacts of Medicaid, Jamila Michener, describes Medicaid recipients as “overwhelmingly poor, disproportionately people of color, and unduly prone to health troubles” [7] and notes that African-American recipients composed 32% of the Medicaid rolls in 2015 while White recipients represented 16% [7]. The extent to which the Medicaid rolls are disproportionately African-American is evident if we consider that the general population in the 2012-2016 time period was estimated to be 73.3% White and just 12.6% African-American or Black [10].

States are given flexibility to expand, change, and improve their Medicaid programs through “demonstration projects” authorized by the federal government as Section 1115 waiver applications [11]. These demonstration projects must be determined by the Secretary of Health and Human Services (HHS) to promote Medicaid’s objectives [11]. During the later years of the Obama Administration, several states such as Arizona and Indiana submitted waiver requests to impose a work requirement for Medicaid eligibility [12]. Conservatives view Medicaid solely as a temporary safety net and believe work requirements will push Medicaid beneficiaries into employment and employer-sponsored health insurance [13]. These work requirements would condition Medicaid eligibility on completing and verifying participation in work or volunteer-related activities for a designated number of hours per week or month [14]. The Centers for Medicare and Medicaid Services (CMS) under the Obama Administration determined that state promotion of employment was not allowed under the Medicaid program or under Medicaid demonstration projects [15].

Despite this prior decision, CMS under the Trump Administration signaled a desire to change how Medicaid Section 1115 waivers are considered and evaluated [16]. On January 11, 2018 CMS announced the Trump Administration’s desire to approve demonstration projects that promote work and community engagement, leading to state submissions of Section 1115 waivers with Medicaid work requirement proposals [16]. To date, Indiana, Arkansas, Maine, New Hampshire, Michigan, Wisconsin, and Arizona’s waiver applications were approved by CMS [17], and Kentucky’s waiver was approved, invalidated by a court decision, and then later re-approved in November 2018 with minimal changes [18]. However, Maine’s newly elected governor, Janet Mills, sent a letter to CMS on January 22, 2019 announcing that Maine will not implement the accepted terms of Maine’s waiver application [19]. Eight more states have currently pending applications [17]. Arkansas, Kentucky, Wisconsin, Michigan, and Arizona require work or community engagement activities for 80 hours per month, while New Hampshire and Indiana require 100 hours per month and up to 20 hours per week, respectively [17]. All of these states except Wisconsin expanded Medicaid under the ACA and all offer varying age exemptions from 50-65 years old and over [17].

Public health advocates and policy makers fear that African-Americans will be disproportionately affected by these work requirements. One reason for this concern is that African-Americans face hiring discrimination, with some employers either refusing to hire them altogether or giving them fewer callbacks than Whites during recruitment [20,21]. For example, a recent study by Quillian et al. found no evidence that hiring discrimination against African-Americans has improved since 1989 [21]. Such discrimination means that African-American Medicaid enrollees would be more likely than other groups to be out of work at any given time and will have more difficulty than others in finding work to meet new requirements. A recent study in Michigan on the state’s Medicaid expansion found that African-Americans had significantly higher odds of being out of work (adjusted odds ratio 1.93, 95% confidence interval 1.50-2.49) when compared to Whites and other races or ethnicities [22].

Losing Medicaid coverage not only potentially harms people’s health by reducing financial access to care, it might also negatively affect household finances because having Medicaid coverage may promote employment and reduce financial strain [23], although evidence on this is mixed [24]. An Ohio evaluation of the state’s Medicaid expansion found 52.1% of employed workers agreed “Medicaid makes it easier to continue working” while 74.8% of unemployed workers agreed “Medicaid makes it easier to look for work” [25]. The argument that African-Americans may be both disparately harmed from work requirements and may have difficulty obtaining new employment without Medicaid coverage demonstrates the disruptive potential of these work requirements. This disruption may have long-term consequences because it has been shown that government programs that are punitive and complex in nature, as Medicaid work requirements may be, politically disempower and demobilize recipients, silencing their voice and ability to advocate for better programs [7]. Indeed, Arkansas’ already implemented work requirement program has complex program language, limited outreach efforts, a confusing enrollment process, and an online enrollment and work reporting system that creates barriers for recipients with low computer literacy or those living in rural or underserved areas with limited access to the internet [26]. This administrative complexity may itself push recipients off of Medicaid.

Work requirements were not a possibility for Medicaid until recently; consequently, there is limited evidence of their effect on Medicaid recipients’ ability to consistently access care. Public health professionals fear that work requirements will reduce access to care by forcing people out of the program or reducing their benefits through sanctions. In this case, sanctions would mean enrollees receive fewer benefits for not meeting program requirements [27]. While the effects of Medicaid work requirements are not yet clear, previous iterations of work requirements under welfare, specifically the 1996 Personal Responsibility and Work Opportunity Reconciliation Act (PRWORA), may provide some insight.

PRWORA, or “welfare reform”, included work requirements that were mandated federally but subject to each state’s welfare policies [28]. Prior to PRWORA, welfare consisted of the Aid to Families with Dependent Children (AFDC) program, which was enacted in the 1930s and provided cash assistance to low-income single mothers [29]. In the 1980s, the Reagan Administration, which popularized the Black “Welfare Queen” stereotypes, became frustrated with work disincentives in AFDC and supported welfare-to-work programs [30]. Similar dissatisfaction on the behalf of states led them to submit waivers to implement work requirements, which 27 states already had when PRWORA passed [30].

PRWORA established the Temporary Assistance for Needy Families (TANF) program, which among other stipulations, established work requirements, imposed time limits for receiving benefits of at most 60 months, added sanctions, and eliminated AFDC and its funding structure [30]. In simple terms, the work requirements in PRWORA dictated that individuals would not be eligible for TANF benefits unless the individual worked an average of 20 hours per week or was covered under an exemption [31]. Before PRWORA was implemented, caseload levels had been stagnant between 1983-1989 but had risen by 27% from 1990-1994 [30]. Levels then declined nationwide by 56.5% from 1994-2000 because of states’ work requirement proposals and PRWORA’s implementation [30]. PRWORA initially had positive economic results, including a 17% increase in the employment of low-income single mothers compared to before welfare reform and child poverty rates dropping for four years [32]. However, 40% of people leaving the welfare program were unemployed, and many of those who gained employment remained in poverty due to low job quality [32].

Several studies suggest that African-Americans may be disproportionately targeted and harmed by welfare restrictions in general and PRWORA work requirements in particular. For example, a study published in the *American Journal of Political Science* (AJPS) found that Whites receiving welfare were sanctioned less than other groups even when all other characteristics are controlled [27]. Additionally, DeParle mentions a 2001 AJPS study by Soss et al. that “tried to quantify the factors that shape state policy. The most important was race: the more blacks [people] on the rolls, the tougher a state chose to be” [33,34].

Recent articles on the new Medicaid work requirement proposals have considered their impact on marginalized groups like disabled individuals but have yet to fully investigate their impact on African-Americans specifically. Sara Rosenbaum, a health policy expert, is cited by *Vox* as warning of “the potential for enormous discrimination, really racial redlining” [35]. The article continues to describe that African-American, urban communities may not benefit from as many exemptions, thereby experiencing a greater burden under the work requirements [36]. These arguments and the scattered reports of welfare reform’s disparate racial impact in sanctioning and retention, indicate the need to study PRWORA more thoroughly as a means of better understanding the potential for work requirements to have a disparate impact on access to Medicaid for African-Americans.

## Objectives

The goal of this paper is to understand the potential for Medicaid work requirements to have a disparate impact on access to health insurance for African-Americans. Our paper aims to systematically review the literature on the racial impact of PRWORA work requirements and highlight aspects of the requirements that seem associated with a disparate racial impact; and discuss the implications of our findings on Medicaid and its recipients.

## Methods

### Search Strategy

The systematic review search query used the ProQuest electronic database and only included scholarly journals in English with a publication date range of January 1991–August 2018. Since work requirements were first proposed for welfare reform, this date range reflects Presidential candidate Bill Clinton’s campaign promise to “end welfare as we know it” in 1991 [33]. The search query read: “Work Requirements AND Personal Responsibility and Work Opportunity Reconciliation Act AND Race OR Barriers to access AND Adults NOT Children.” The rationale for each included search term is as follows:

### Work Requirements

The focus of this study was specifically on the work requirement proposals included in the recent Medicaid Section 1115 waiver applications, which is why this term was first in the search query. “Work Requirements” was chosen as a search term instead of “welfare reform” since there are many parts to PRWORA and this search term would filter results more towards the study’s focus.

### Personal Responsibility and Work Opportunity Reconciliation Act

This term was included in the query to ensure the search results were curated towards PRWORA.

### Race

“Race” was included since the main purpose of this investigation was to understand if Medicaid work requirements would have an impact on people of color, and specifically on African-Americans. Since the Office of Management and Budget’s Directive No. 15 stipulates that race and ethnicity data should be collected separately, the potential exists for studies on welfare to solely focus on race; therefore, this search query solely mentioned race [37].

### Barriers to Access

The “Barriers to Access” term was included in the query because the overall outcome of interest in the studies reviewed was a difference in who was denied access to the welfare program. The prefix to the term in the search query was “OR” so search results focused on race or on barriers to access would be selected in the search results.

### Adults

The “Adults” search term was included in the query because work requirements do not often apply to individuals under the age of 18 and this term limited articles towards the population of interest.

### Children

An initial test search query included multiple articles focused on children; so the search term “NOT children” was included to ensure articles on adults were selected instead of children and the Children’s Health Insurance Program.

### Selection

The review strove to identify quantitative studies specifically focused on welfare *sanctions* and *caseload movements* as these outcomes have been identified as indicators of welfare racism [38]. We defined “caseload movements” as a change in recipient status that is different from prior to when TANF was enacted or that is different compared to other groups on TANF. Selected articles also had to focus on TANF, adults 18 years old and older, and discuss or present data on the demonstrable impact of these outcomes on African-Americans. Articles that did not meet these criteria were labeled ineligible and were excluded. Documentation of articles included or excluded as well as extent of screening is provided in the PRISMA tree below (see Figure 1).

**Figure 1:**
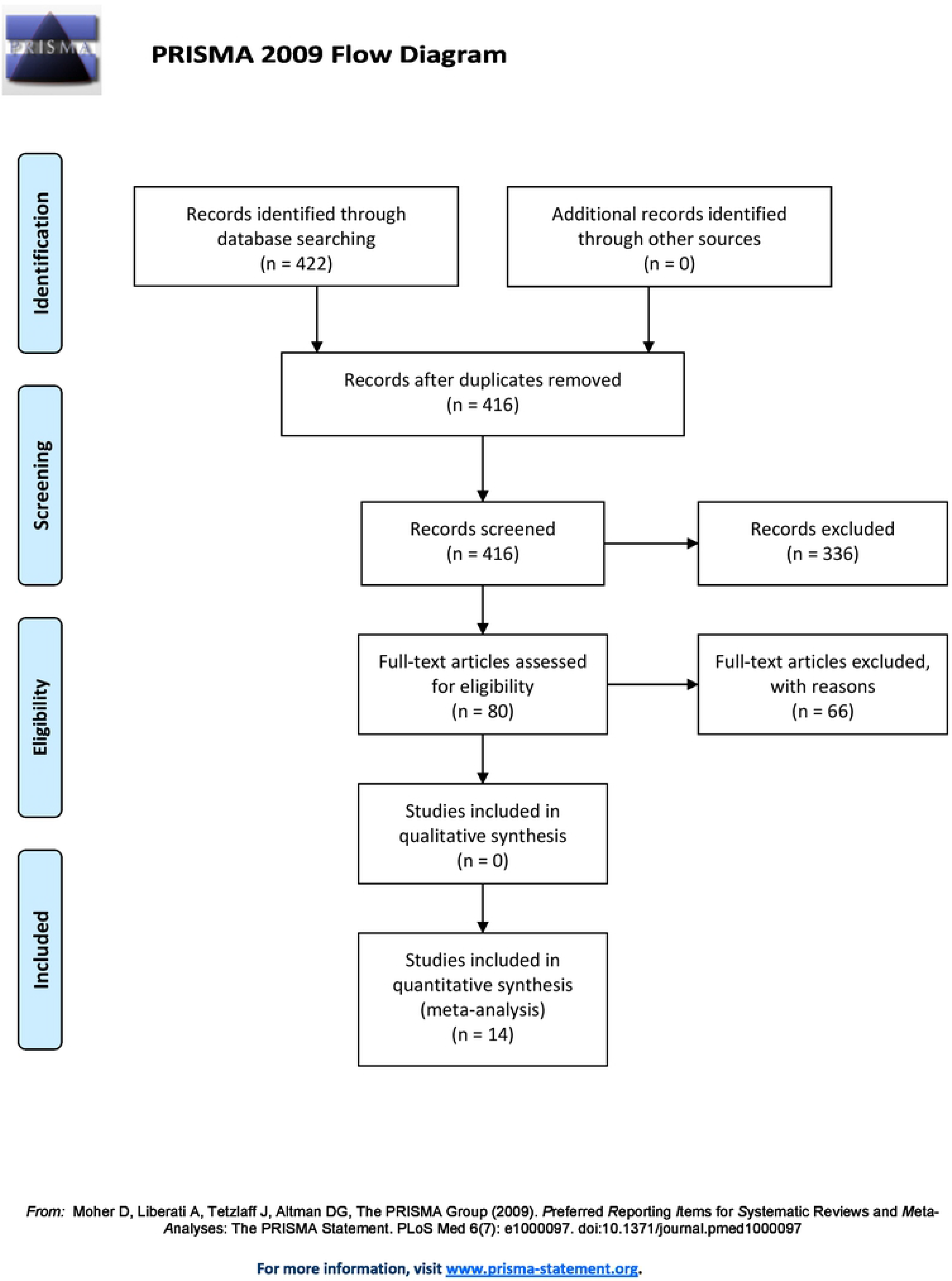
PRISMA Flow Diagram

### Critical Appraisal and Data Extraction

Selected articles were grouped by their respective outcomes and summarized in a table by their study design, location, outcome studied, population studied, population size, covariates, and relevant findings with an appreciable racial impact on African-Americans being the main focus. An appreciable racial impact was defined as African-Americans experiencing the outcomes of interest more or less frequently or severely than Whites or other races. Critical appraisal of selected articles was performed using an adapted form of the National Institutes of Health’s (NIH) Quality Assessment Tool for Observational Cohort and Cross-Sectional Studies [39]. Other than this adapted quality assessment tool, no previously developed protocol was used for this systematic review.

## Results

This search strategy identified 422 scholarly journal studies with 14 of those studies selected for inclusion in the final analysis. Other articles types that were eliminated included dissertations and theses, reports, and working papers. Six of the 422 studies were duplicates and after 336 were excluded based on the studies’ abstract or introduction, 66 of the 80 studies read in full were excluded either due to a wrong outcome, being a qualitative study, not focusing on African-Americans, TANF, or adults ages 18 and over, and not presenting significant results. The included items were all quantitative studies with a time frame of 1990-2006. Studies used secondary data analysis as well as cohort, cross-sectional, and longitudinal designs to investigate welfare reform and the effect it had on its recipients. These studies analyzed TANF from multiple levels of government within the United States, including the federal level, the state level (Wisconsin, Michigan, Maryland), the municipal level (New York City, Boston, Chicago, San Antonio, and 20 unspecified cities in 15 different states) and the county level (an unspecified Michigan county). Population groups studied included women, single mothers, mothers in general, parents in general, adults, caregivers in low-income families, TANF recipients, and households. Study details and designs are listed in Table 1 and grouped into the two outcomes of interest: Caseload Movements and Sanctions. Table 2 lists quality designations for each study.

**Table 1:**
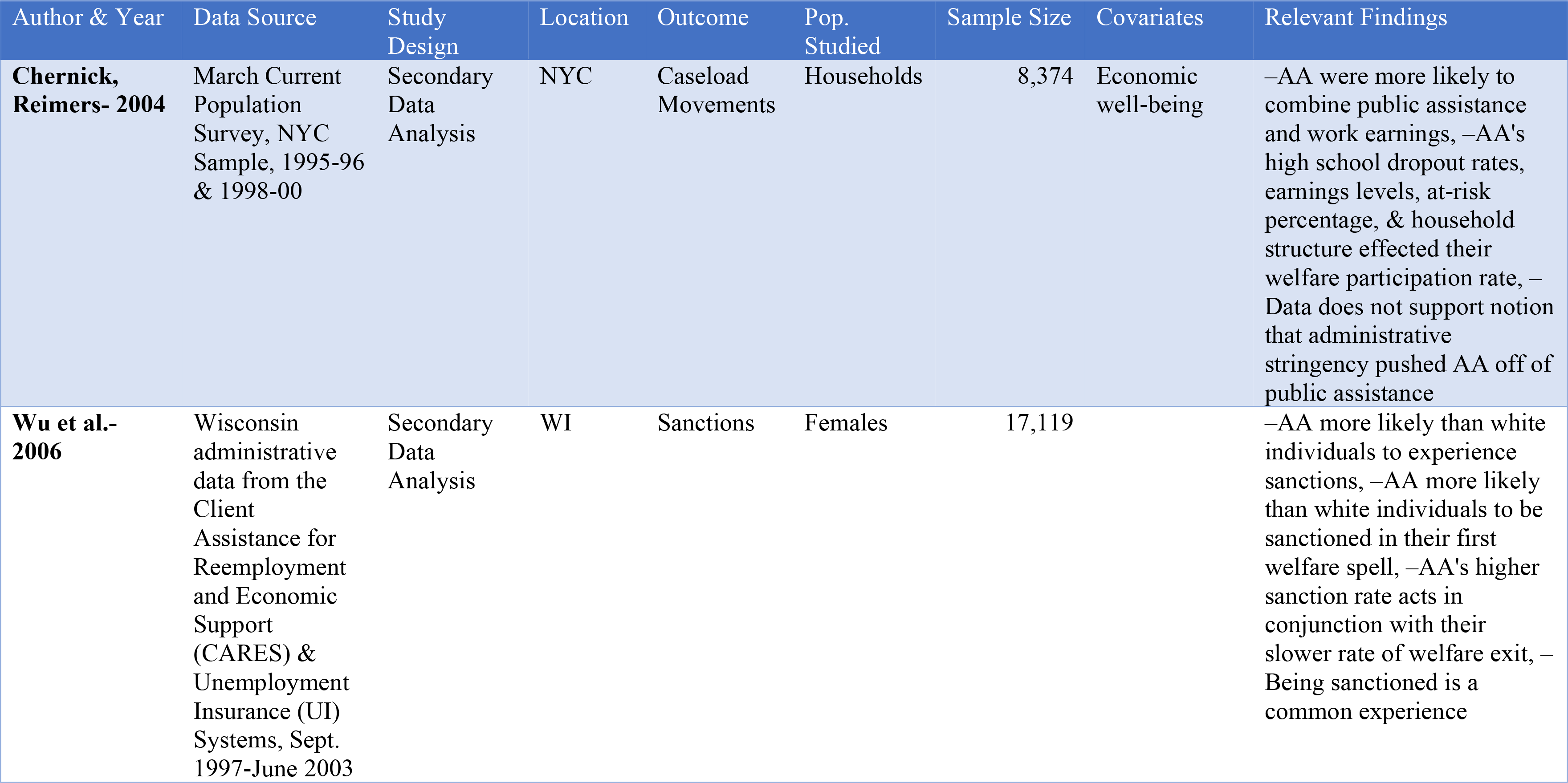
Selected Study Information

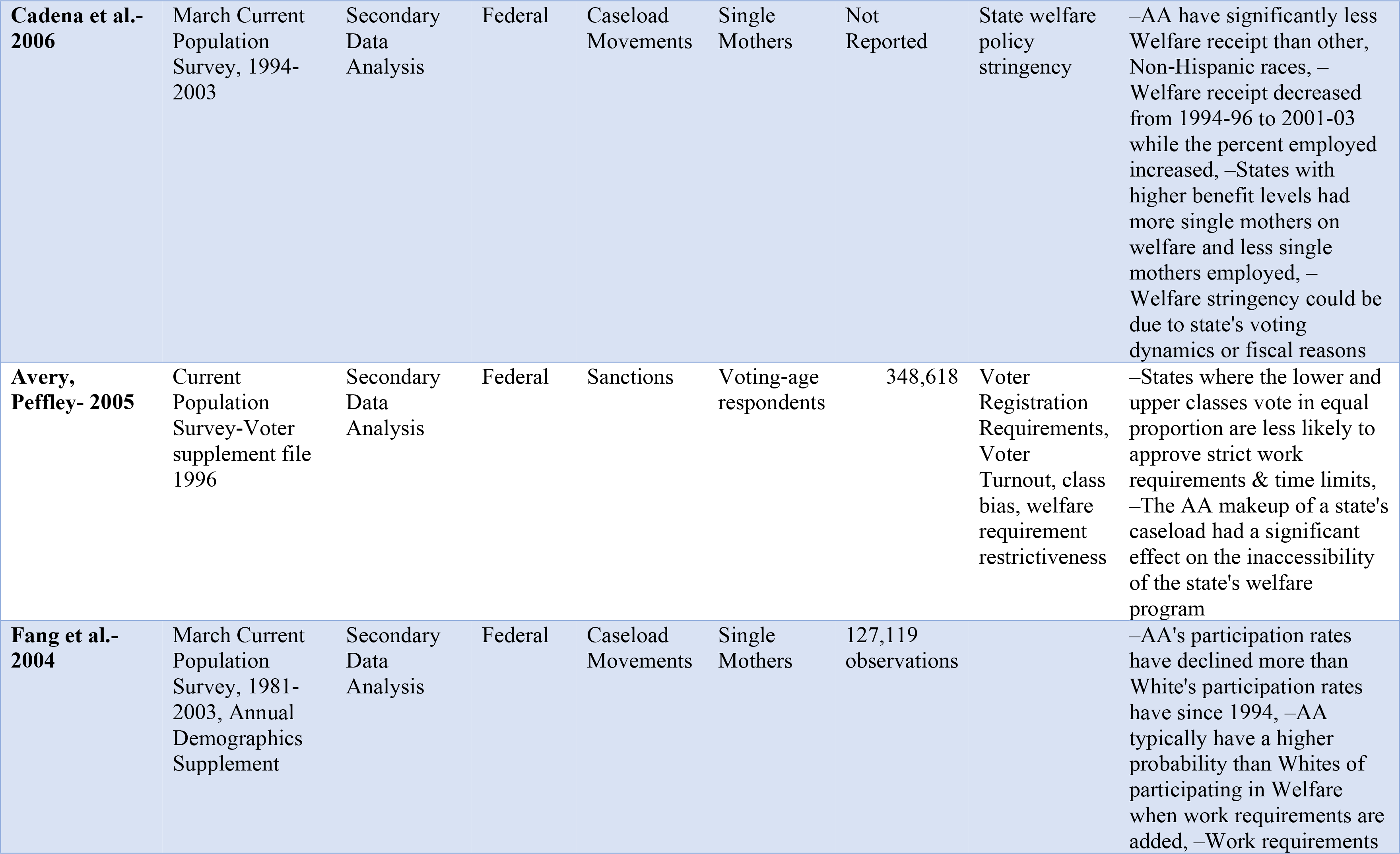

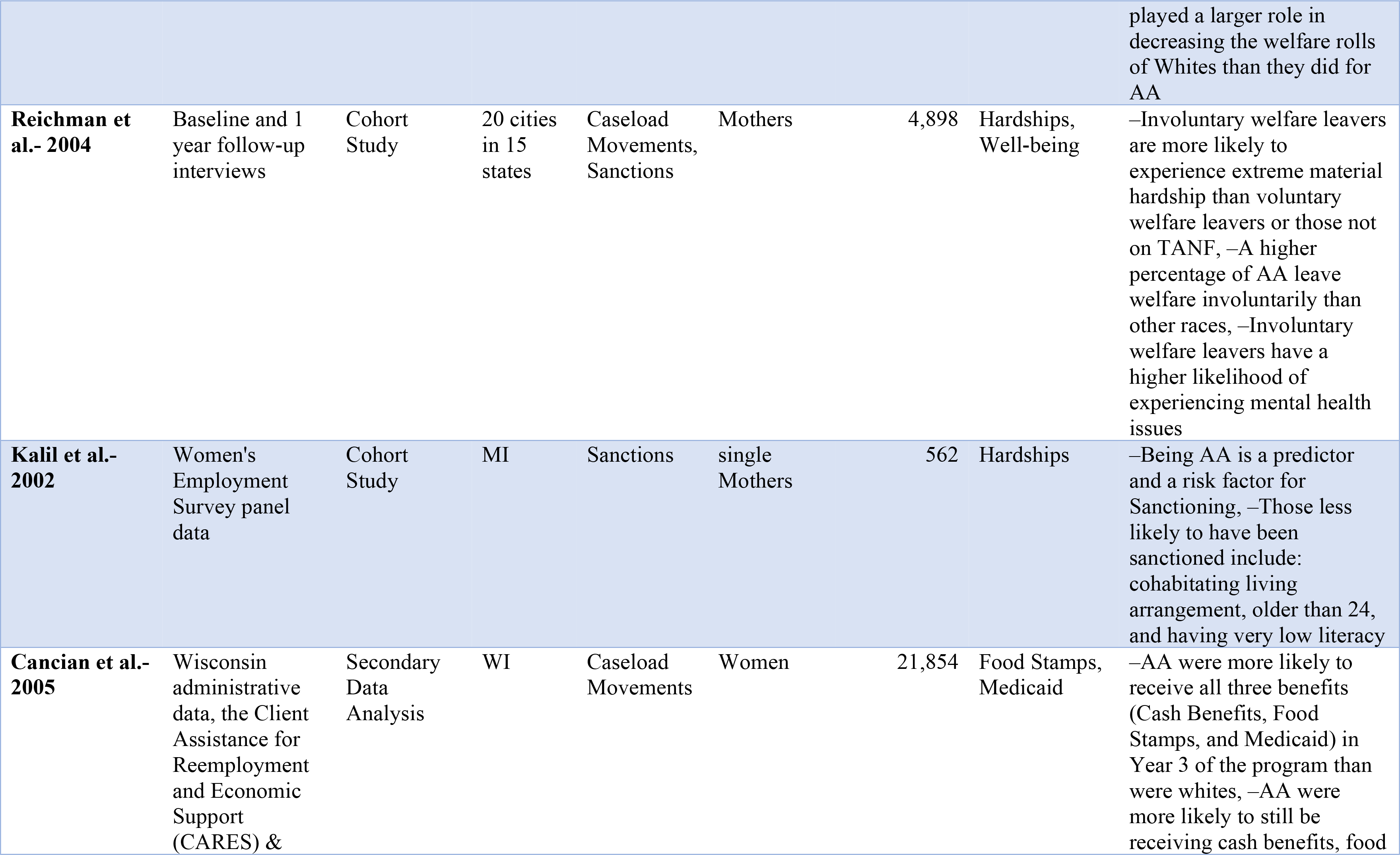

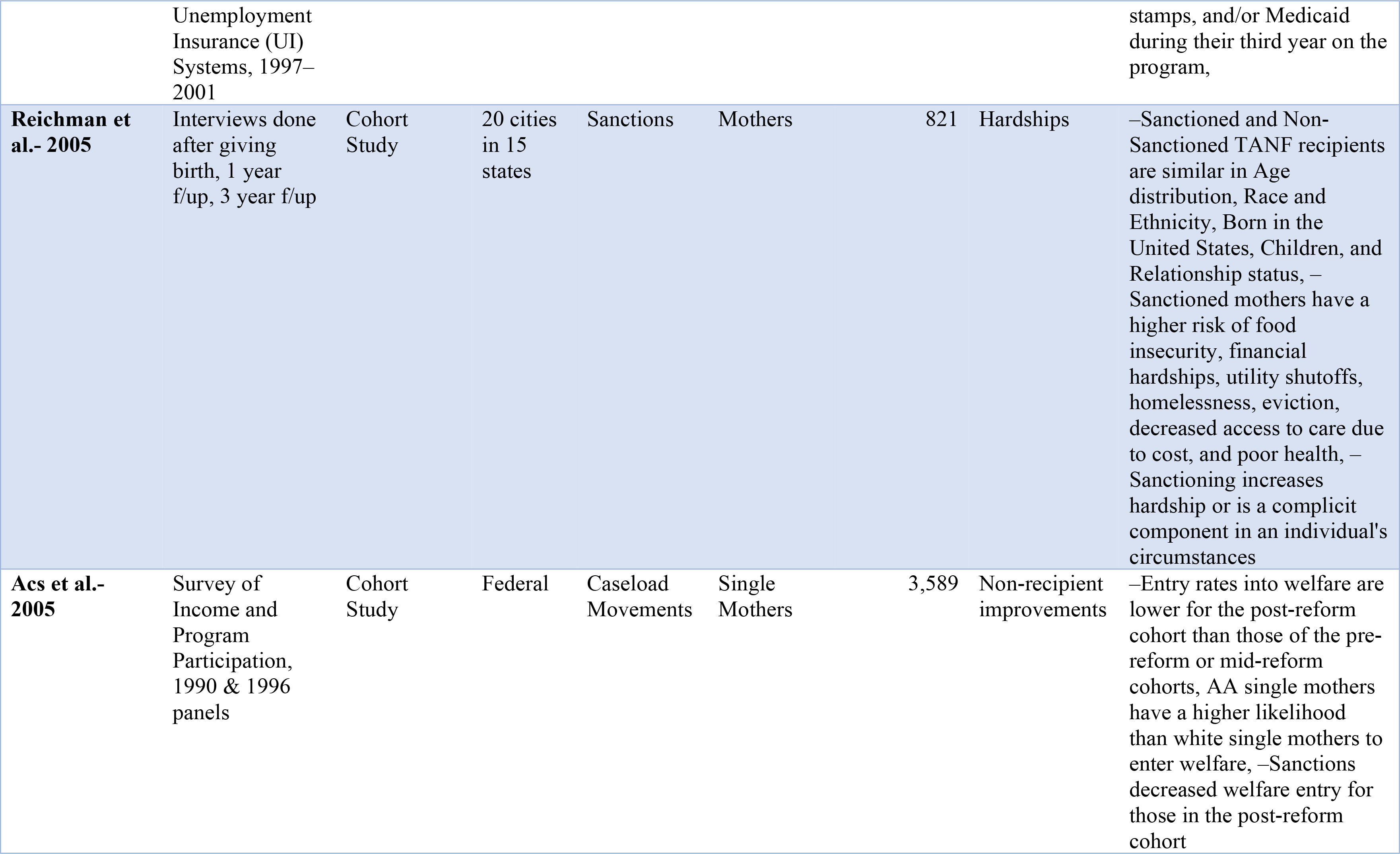

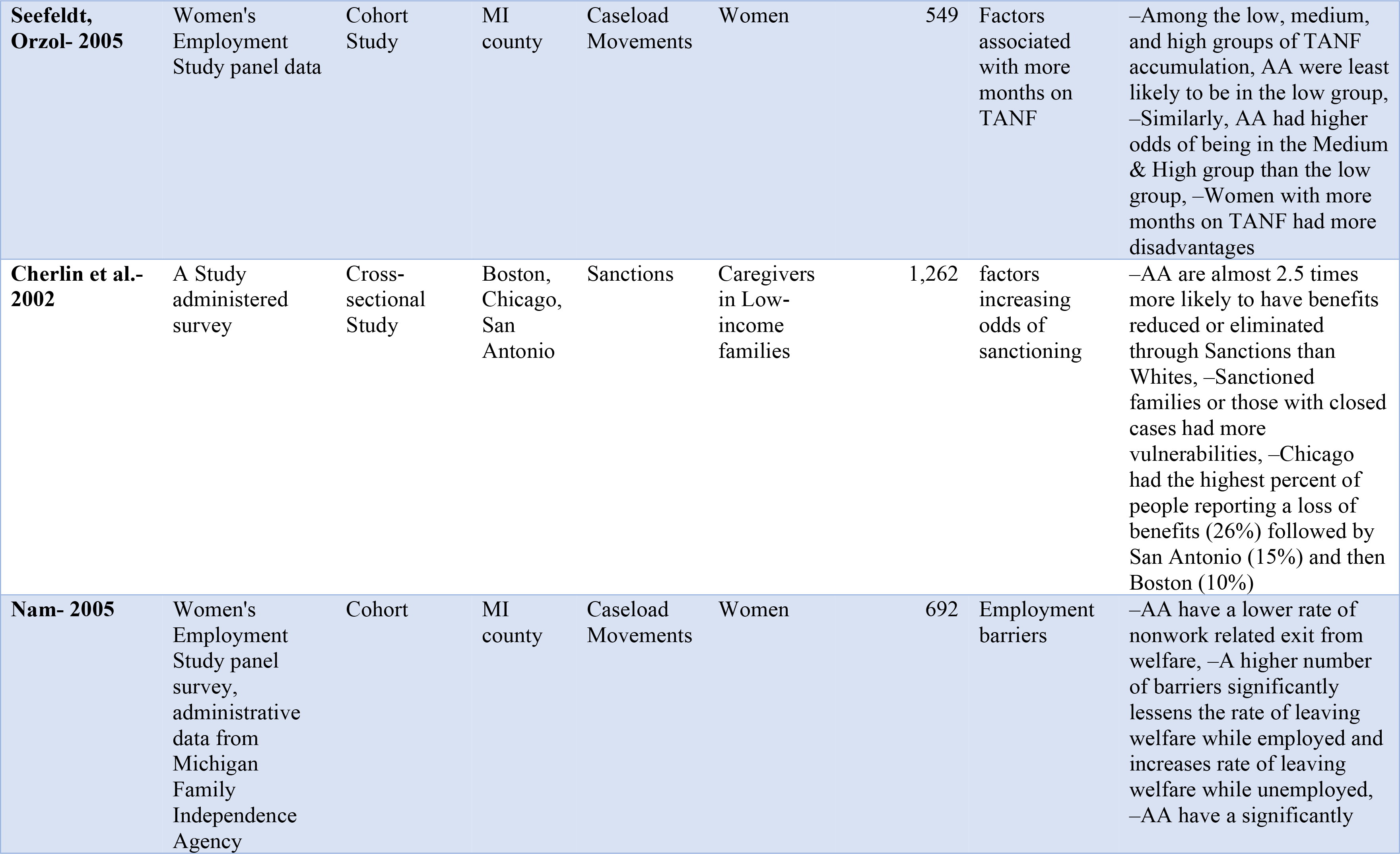

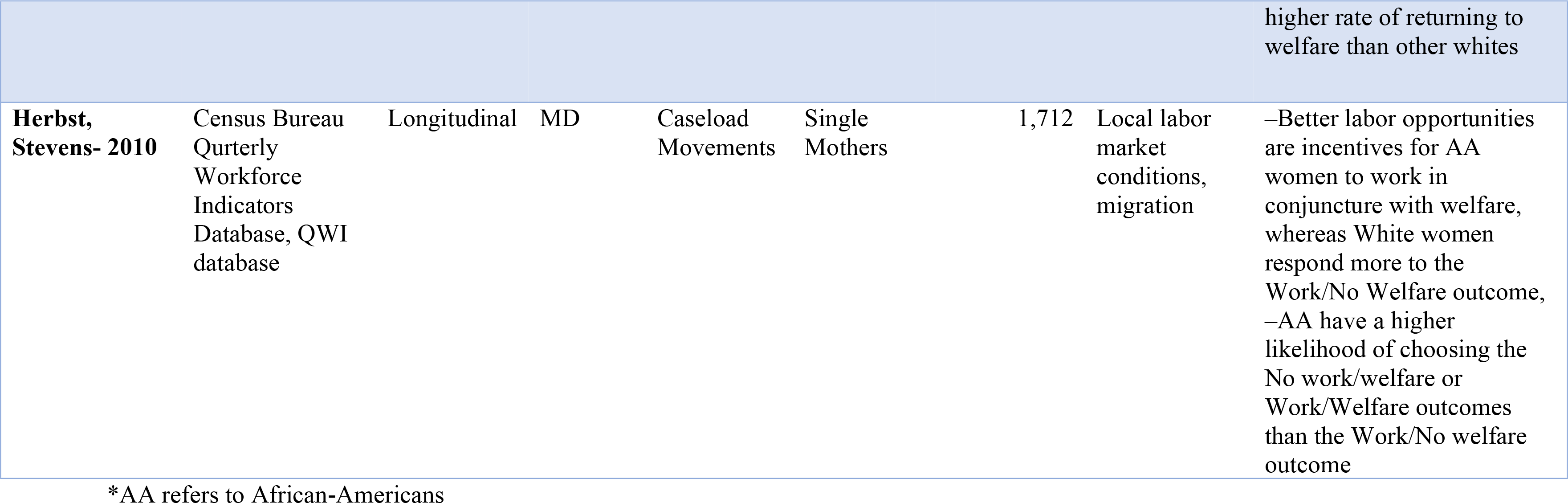

**Table 2:**
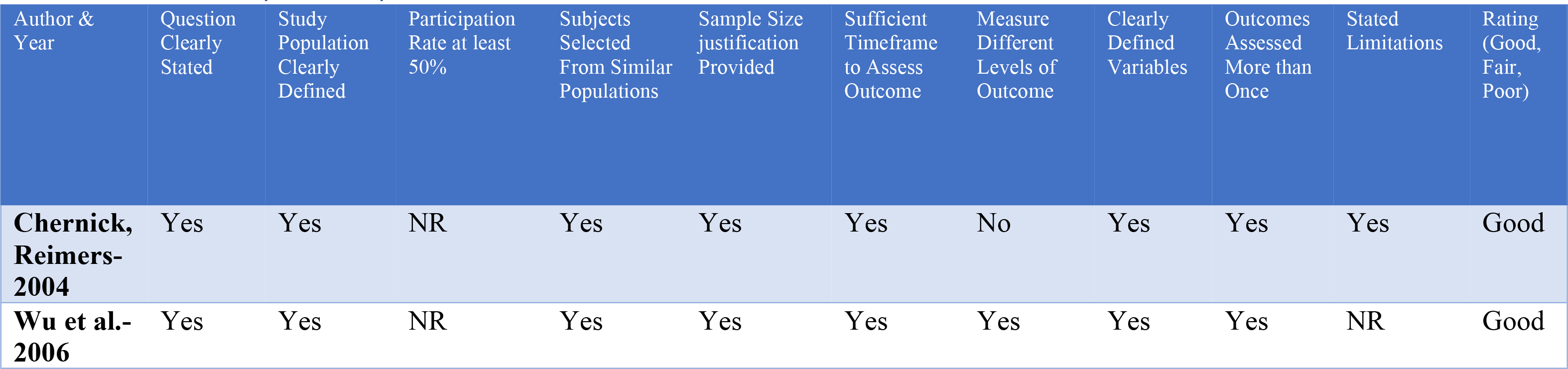
Study Quality Assessment

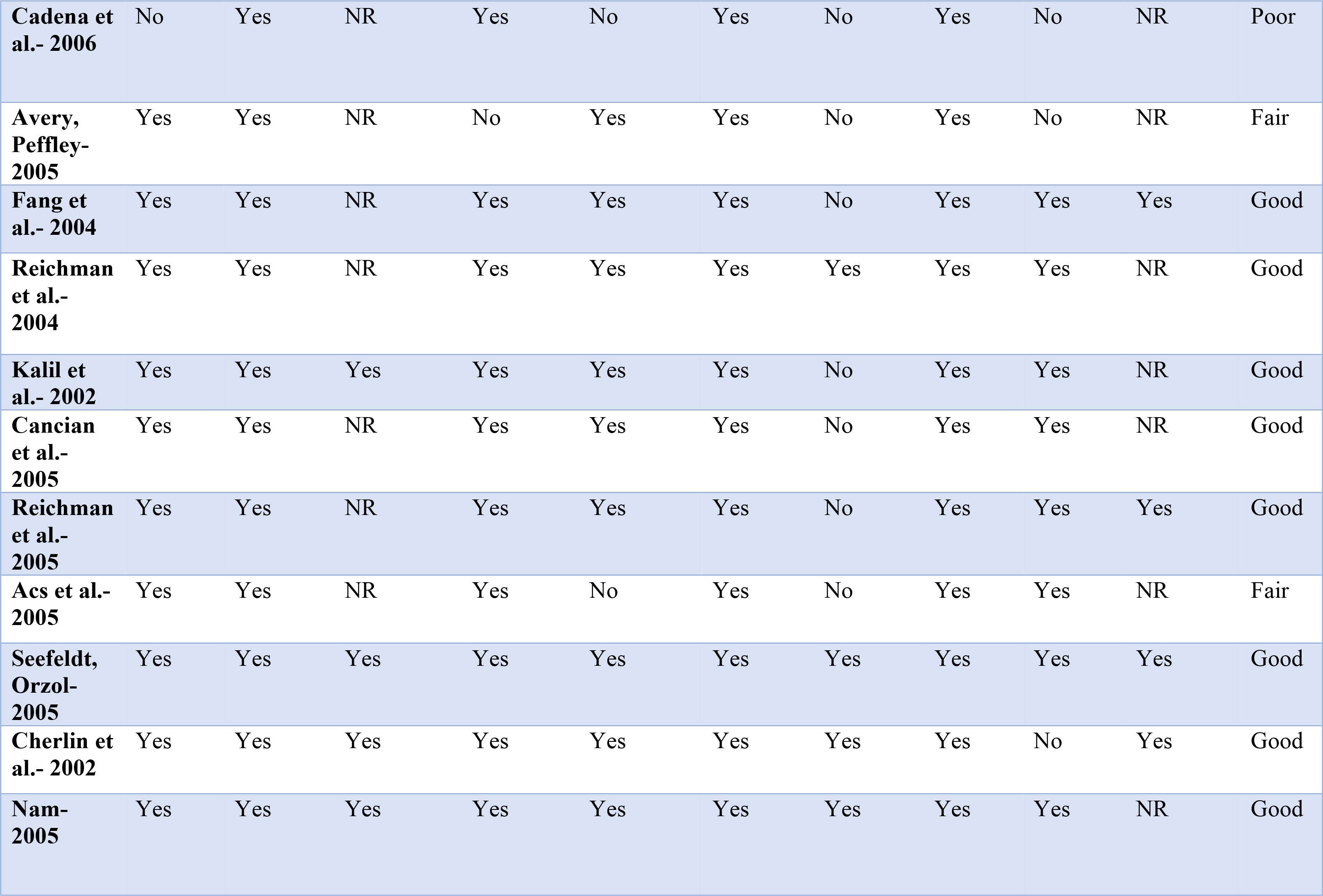

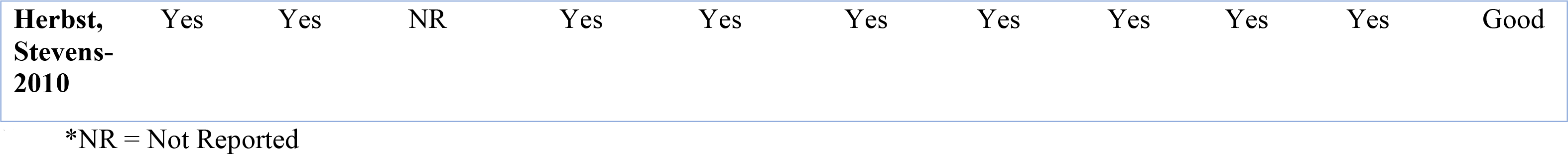

We have mixed findings on the impact of work requirements on African-Americans’ access to welfare programs. Overall, the studies reviewed did not find that African-Americans were removed from TANF at higher rates than Whites; instead, most studies examining caseload movements found that African-Americans entered TANF at higher rates and remained on TANF longer than Whites. However, the studies reviewed find that under work requirements, African-Americans experienced sanctions more frequently and stringently than Whites.

### Caseload Movements

Of the selected studies, eight analyzed caseload movements by examining movements on and off welfare, retention disparities, and modeled predicted movements by race. Two of the three studies looking at the populations moving on and off welfare generally found that African-Americans entered welfare at higher rates and exited welfare at lower and slower rates than whites and other races. For instance, Acs et al. compared pre-TANF and post-TANF periods and found that African-American single mothers significantly increased their rates of entering welfare while White’s rates decreased [40]. Additionally, Chernick and Reimers found a significantly different drop in the welfare rolls during TANF with African-American households’ enrollment decreasing by 10% while White and Hispanic households decreasing by 32% and 42% respectively [41]. Importantly, African-American households moved from the “welfare receipt only” group to the group receiving both welfare benefits and work earnings at a higher rate than other races, although, it is not clear if this difference was statistically significant [41]. Contrary to the previous two studies, Cadena et al., the third study focusing on movements on and off welfare, found that when state welfare policy increased in stringency by one standard deviation, African-American single mothers had a significant 5.7% decrease in their welfare benefit receipt, which did not occur with other, non-Hispanic races [42].

Retention disparities measures the lack of caseload movements. Four of the selected studies focused on retention and found that compared to Whites, African-Americans were significantly less likely to become ineligible for TANF [40]; significantly more likely to still be receiving welfare in their third year [43]; had significantly higher rates of returning to welfare [44]; and were significantly more likely to receive welfare between 20 and 60 months while Whites were more likely to receive welfare for less than 20 months [45].

Finally, two studies used models to predict the likelihood of caseload movement and generally found that African-Americans were predicted to be more likely to participate in welfare or combine welfare benefits and work earnings than Whites. Fang et al. predicted African-Americans would have a higher probability of participating in welfare when a work requirement was added and that work requirements, when implemented, would account for a greater percentage of caseload decline for Whites than African-Americans [46]. The other study, by Herbst and Stevens, found African-Americans would be significantly more likely to combine work earnings and welfare benefits while Whites would be significantly more likely to choose work without welfare benefits [47].

### Sanctions

Six of the 14 studies selected for analysis focused on the sanctioning frequency and severity African-Americans experienced under TANF. Four of those studies found that African-Americans experience more frequent sanctions than Whites. Specifically, when compared to Whites, studies found that African-Americans were significantly more likely to experience sanctions during their first welfare spell, in part due to their significantly slower rate of exit [48]; were involuntary TANF leavers due to sanctions or time limits at a significantly greater percentage (57%) than White mothers (11%) [49]; and were 1.73 times more likely to experience sanctions [50]. Additionally, one study found that African-American caregivers in low-income families had an odds ratio of 2.45 of having their TANF benefits reduced or eliminated, although the authors note this is not statistically significant, partly due to the low number of White caregivers sampled [51]. Reichman et al. also found no significant difference between sanctioned and non-sanctioned mothers in terms of race and ethnicity among other demographic factors [52].

Two of the selected studies focused on sanction severity, finding that African-Americans experienced more stringent sanctions and states would become more stringent when higher percentages of their caseloads were Black. Specifically, Wu et al. found that when compared to White mothers, African-American mothers had a significantly higher likelihood of receiving partial and full sanctions during their first welfare spell [48]. Avery and Peffley predicted that states with higher percentages of Black individuals on their caseloads would have an increased probability of more stringent sanctions, by a significant difference of 36%, when compared to states with lower percentages of their caseload being Black [53].

## Discussion

This study set out to review the literature on racial disparities in TANF access due to work requirements, specifically the impact on caseload movements and sanctioning. We reviewed 14 studies from a search of 1,672 studies and found that with the introduction of work requirements, African Americans were not removed from TANF at higher rates than Whites but rather entered TANF at higher rates and remained on TANF longer than Whites. However, the studies reviewed also found that African Americans experienced sanctions under TANF more frequently and stringently than Whites.

The caseload movement results are surprising, as prior to this systematic review, we would have hypothesized that African-Americans would be disproportionately pushed off the TANF rolls when compared to other races due to hiring discrimination, higher odds of unemployment among African-Americans, and state welfare policy toughness. The results point to the persistent disparities in wealth and income, driven by profound historical and contemporary racial oppression, that results in differential need for welfare among different racial and ethnic groups in the United States. The results from Herbst and Stevens as well as Chernick and Reimers showing that African-Americans would be significantly more likely than Whites to combine both work earnings and welfare benefits lends credence to this idea as they suggest that when they were working, African-American TANF beneficiaries may have been less likely to earn enough to reduce the need for supplemental welfare assistance. Work requirements and TANF undermined the stronger safety net program that was AFDC; similarly, work requirements when added to Medicaid threaten to undermine this essential public health insurance program.

It is not clear what the implications of these findings are for Medicaid work requirements. It could be that Medicaid caseload movements will be more pronounced for African Americans compared to that seen under TANF because of the different benefits being offered. Benefits under TANF may be more tangible, allowing TANF recipients to survive until the next installment of benefits, and provide motivation for recipients to continue navigating burdensome work and reporting requirements. However, Medicaid benefits– health care access to varying extents– may be less tangible as recipients may postpone health care visits until truly necessary and dire when faced with access barriers [54]. In contrast to TANF, this may provide less motivation to navigate burdensome requirements. The recent Department of Homeland Security’s proposed public charge rule change already defines benefits differently by labeling their ability to be monetized; TANF can be while benefits like Medicaid are classified as “cannot be monetized” [55]. The question of how differential benefits may effect motivation and caseload movements represents a limitation of this study and future research should investigate this question once more data from Medicaid work requirement programs becomes available.

The significant results on sanctions represent our main assessment of how the history of work requirements for welfare inform our understanding of the implications of work requirements for Medicaid. The studies reviewed found that African-Americans were disproportionately harmed by sanctions under TANF. Only one study from Reichman, Teitler, and Curtis found no significant difference between sanctioned and non-sanctioned mothers in terms of race. However, Reichman et al. acknowledge study limitations that include not differentiating between sanctions that reduce and terminate benefits and that their follow-up data was self-reported, which, given that some women did not know their sanction status, they believe this might have underestimated sanction effects.

Many of the Section 1115 waiver applications for Medicaid work requirements mention lockout periods, in which recipients who remain unemployed or do not correctly follow work and community engagement reporting guidelines may be locked out of coverage for a set number of months. This is where TANF sanctions serve as the best guide for assessing Medicaid. If African-Americans are disproportionately subject to lockout periods, as the TANF sanction results suggest, this may result in fewer African-Americans receiving health care services, more preventable deaths and diseases going untreated in the African-American community, a higher burden placed on emergency departments, and more expensive medical bills being paid by African-American patients out-of-pocket or passed along to state governments.

While this study focused exclusively on the potential impact of Medicaid work requirements on African-Americans, other races will also be disproportionately affected by these requirements. A number of the welfare studies also reported on caseload movements and sanctions’ impact on Hispanic people. Future research should assess the potential for Medicaid work requirements to harm Hispanic communities. We must also note a limitation of the eligible welfare studies is that races such as Asian-Americans were not represented in the results.

### Limitations

Limitations of this study include the fact that assessing current legislation based on a law signed over 20 years ago is inherently challenging due to changing socio-political contexts. In addition, state flexibility in both welfare and Medicaid makes it difficult to compare data across studies and programs. The eligible welfare studies that we reviewed mostly focused on women whereas Medicaid work requirements may more equally effect multiple genders. This limits the generalizability of our findings. Finally, we note that just one researcher conducted the search, screening, and review process. One notable strength is the long time period allowed for eligible studies, which given state flexibility, allowed us to include studies related to a change in a jurisdiction’s welfare policy that happened later than the immediate years after 1996.

### Recommendations

We have identified a number of policies that might mitigate the negative impact of Medicaid work requirements. First, states could enact policies that prohibit work activities and community engagement from being conditions of Medicaid eligibility. For example, California passed SB 1108 in September 2018 with these stipulations [56]. States can also strengthen Medicaid via Section 1115 waivers not focused on work requirements. This could include expanding the populations covered under Medicaid, increasing and improving services covered, such as mental health or dental services, and solving Medicaid’s transferability issues. Additionally, states could pass evidence-based policies that combat hiring discrimination or pass “Ban the Box” policies to benefit job applicants with criminal histories. As part of their approval checklist, CMS could scrutinize states’ policies on this matter when they consider future Section 1115 waivers involving work requirements. Finally, states could, at a minimum, decrease the burdensome reporting requirements that exist for recipients of work requirement programs. Arkansas Medicaid recipients, for example, should not lose their health insurance coverage simply because they have limited access to internet services or the technological knowledge necessary to navigate the reporting website. With 18,164 Arkansas Medicaid recipients losing coverage for failing to meet work requirements since September 2018, these policy changes are critical [57].

## Conclusion

Our systematic review of TANF literature found that after the introduction of work requirements, African-Americans entered TANF more than Whites, remained on TANF longer, but experienced sanctions more frequently and severely than Whites. These results suggest that while Medicaid work requirements might not disproportionately reduce enrollment rates for African-Americans, it may reduce their access to care through sanctioning such as lockout periods. At a minimum, the caseload movement results illustrate the need for strong safety net programs, which are not the intent of work requirements for Medicaid. Recommended policy changes include prohibiting work activities as a condition of Medicaid coverage, strengthening Medicaid apart from work requirements, fighting hiring discrimination, and easing burdensome reporting requirements. Once more data on the impact of these work requirements becomes available, future research needs to investigate the racial impact of caseload movements and lockout periods and how they contribute to disparate health outcomes. Given that the National Medical Association, composed solely of African-American physicians, was the only physician group to support Medicaid when it was proposed in 1965, we owe those physicians and the African-American community a thoughtful and rigorous analysis of work requirements on this vital program. Future research should also seek to understand if a difference in benefits offered from safety net programs is significant enough to impact motivation to remain on programs despite burdensome and complex requirements. Such research may improve our ability to correctly assess the impact of future programs based on prior ones.

## Acknowledgements

We thank the faculty and staff at Touro University California, especially Dr. Deirdra Wilson, Dr. Carly Strouse, and Dr. Jennifer Abueg, who were critical advisors on this paper. Additionally, we thank Tory Cross for the time and efforts she spent editing and providing crucial feedback throughout the researching and writing process. As this paper was completed as a Master’s thesis, there are no funding sources to declare.

## Supporting Information Captions

S1 Checklist: PRISMA 2009 Checklist

